# Replication-deficient Adenovirus 5 Serotypes Induce Type I Interferon and enhance BCG-mediated Immune Response in Co-infected Murine Macrophages

**DOI:** 10.64898/2026.06.24.734229

**Authors:** Joseph A. Vecchio, Jeffrey S. Schorey

**Affiliations:** Department of Biological Sciences, University of Notre Dame, Notre Dame, IN, USA; The Berthiaume Institute for Precision Health, University of Notre Dame, Notre Dame, IN 46556, USA; Eck Institute for Global Health, University of Notre Dame, Notre Dame, IN 46556, USA

## Abstract

Tuberculosis (TB) remains a leading global cause of infectious mortality due, in part, to the limited efficacy of the *Mycobacterium bovis* BCG vaccine against pulmonary TB. Previous studies in mice have shown that stimulating type I interferon (IFN) signaling during BCG vaccination can bolster protection against *Mycobacterium tuberculosis*, yet clinically feasible delivery strategies for this approach are lacking. Adenoviral vectors, which induce potent type I IFN responses and are utilized in approved vaccine platforms, represent a promising adjuvant strategy. To evaluate the host immune response to this combination, bone marrow-derived murine macrophages were co-infected with replication-deficient adenovirus and BCG. Adenovirus-infected macrophages elicited a robust type I IFN response via the cGAS/STING pathway. Compared to BCG infection alone, co-infected macrophages exhibited additive expression of genes with known host-protective roles against *M. tuberculosis*. Conversely, co-infection with BCG suppressed adenovirus-induced type I IFN signaling and diminished the production of IFN-stimulated genes compared to adenovirus infection alone. Together, these findings reveal a complex regulatory interplay during adenovirus and BCG co-infection. While BCG partially restricts adenoviral IFN induction, the co-infection still drives an enhanced host-protective gene profile, suggesting that adenoviral vectors could serve as a viable platform to modulate innate immunity and improve BCG vaccine efficacy.

**IMPORTANCE:** Tuberculosis (TB) remains the leading cause of death by a single infectious organism with approximately 1.25 million deaths annually. *M. bovis* BCG remains the only approved vaccine for TB; however, its efficacy against the contagious and most common pulmonary form of the disease is limited. There have been numerous attempts to improve BCG efficacy, but these approaches have not resulted in any clinically approved vaccine. We propose that BCG combined with a replication-deficient adenovirus presents a way to bolster vaccine-conferred protection as the combination may elicit a robust innate immune response and drive a more protective T cell response. Moreover, BCG and replication-deficient adenoviruses have well-assessed safety profiles and decades of studies regarding their use in patients. The significance of our work is in leveraging their complementary immunology to function as a combined vaccine platform. This approach presents a novel and clinically feasible approach to improve the BCG vaccine.

## INTRODUCTION

Tuberculosis (TB) is the leading cause of death from a single infectious agent globally accounting for approximately 1.25 million deaths annually. There is only one WHO-approved vaccine developed over a century ago. (1). The vaccine, which is an attenuated strain of *Mycobacterium bovis* (*M. bovis* BCG), is effective against severe, disseminated forms of disease in young children, such as meningeal disease and TB meningitis (2–4). However, it lacks efficacy against the pulmonary form of the disease, which is the form responsible for person-to-person transmission (5, 6). Hence, there is a major need and opportunity for improving the efficacy of the BCG vaccine.

Type-I interferons (IFNs) are critical drivers of innate immune responses with well-established protective roles during viral infection. Viral components bind pattern recognition receptors present on the cell membrane, inside endosomal compartments, or in the host cytosol to trigger signaling pathways leading to the production of type-I IFNs via transcription factors such as IRF3 and IRF7 (7). Type-I IFNs are secreted out of the cell and bind to their receptor IFNAR in an autocrine or paracrine manner to activate JAK-STAT signaling. Activation of the JAK-STAT pathway results in the transcription of hundreds of interferon-stimulated genes (ISG) that work to inhibit viral replication (7, 8).

The role of type-I IFN signaling in mycobacterial infections are more complex and less understood (9–12). While certain mycobacterial strains, such as *M. tuberculosis* (MTB) and *M. smegmatis* (MSM), have been noted for their ability to induce higher levels of type-I IFN (13, 14), BCG generally elicits a weakened type-I IFN response due, in part, to a deletion of the ESX-1 secretion system (15, 16). The ESX-1 secretion system enables other species like MTB to gain access to the host cytosol and activate cytosolic pattern-recognition receptors (14, 17–19). Type-I IFNs, specifically IFN-β, have been associated with MTB pathogenesis, and their deleterious roles during infection have been well-characterized in both mice (20–22) and humans (23–26). Paradoxically, in the context of BCG vaccination, numerous animal studies support the importance of type-I IFNs in providing improved protection against a subsequent MTB infection. There have been several approaches to bolster type-I IFN signaling elicited by the BCG vaccine through the addition of exogenous cytokine (27, 28) or the generation of recombinant strains. These recombinant strains conferred BCG access to the cytosol (29–31) or overexpressed agonists that activate the type-I IFN signaling pathways or antagonists to type-I IFN negative regulators (32–34). These approaches demonstrate that type-I IFN in the context of BCG vaccination can lead to increases in host-protective cytokines, enhance polyfunctional T-cell populations, reduce MTB bacterial burden, and improve lung pathology following MTB infection. While these approaches enhance the protective efficacy of the BCG vaccine, several limitations preclude their translation into clinical practice. Using exogenous cytokines as vaccine adjuvants is particularly challenging in human patients due to systemic toxicity and rapid clearance in-vivo resulting from their short half-lives. Despite efforts to lower virulence, administering a recombinant BCG strain would raise biosafety concerns. Therefore, other mechanisms are needed to induce type-I IFN, in the context of a BCG vaccination, that can be more clinically applicable.

Replication-deficient adenoviruses (RdAds) are highly effective platforms for gene therapy and vaccine development against cancer and infectious diseases (35). These non-enveloped, linear dsDNA viruses have a genome of roughly 30 kilobases and are rendered replication-incompetent by deleting the E1 and E3 genomic regions (35). RdAds represent a strategy for enhancing BCG-mediated protection due to their established safety profiles, current clinical applications, and ability to induce type-I IFN signaling (35–39). We hypothesized that a simultaneous administration of RdAds with BCG will leverage their complementary immunology and bolster the BCG protective responses by relying on RdAds to bolster type-I IFN signaling while BCG provides the antigen-specific responses.

However, it remains unknown how combining BCG and adenovirus modulates the macrophage response, including the production of type-I interferon. To begin addressing this question we infected macrophages with BCG, RdAds, and in combination. We found that the RdAds induced a robust IFN-β response that was STING-dependent. We also found that co-infection modulated the expression of immune regulatory genes when compared to BCG or adenovirus infection alone. The data highlight the complex interplay between adenovirus and BCG but also suggest that the addition of an adenovirus has the potential to enhance BCG-induced protection through targeted modulation of innate immunity.

## RESULTS

### Macrophages co-infected with BCG and RdAds drive MyD88-dependent increases in eGFP transgene expression

To characterize infection rates, samples were imaged using confocal microscopy at 24 hpi to observe BCG and Ad5 uptake via DsRed and eGFP fluorescent reporters, respectively. Upon co-infection, bone marrow-derived macrophages (BMDMs) exhibited higher levels of eGFP expression compared to Ad5 alone (**Figure 1A**). It has been previously reported that the CMV promoter, which is the promoter driving GFP expression, can be influenced by transcription factor binding sites present on the CMV promoter. To test whether the increased GFP expression in co-infected BMDMs was due to known BCG-induced signaling pathways, we utilized MyD88 knockout BMDMs. As predicted, these MyD88^-/-^ BMDMs show significantly diminished *Tnf* responses upon BCG infection compared to WT BMDMs (**Figure S1**). Cells co-infected with BCG and Ad5 also exhibited significantly decreased eGFP expression in the MyD88 knockout compared to WT co-infected BMDMs (**Figure 1B-C**). This trend was also observed across both adenovirus serotypes, Ad5 and Ad5F35, at the transcriptional level (**Figure 1C**). This difference between WT and MyD88^-/-^BMDMs was not observed for Ad5 only infected cells, indicating this was a BCG-mediated effect (**Figure 1B**).

**Figure 1.**
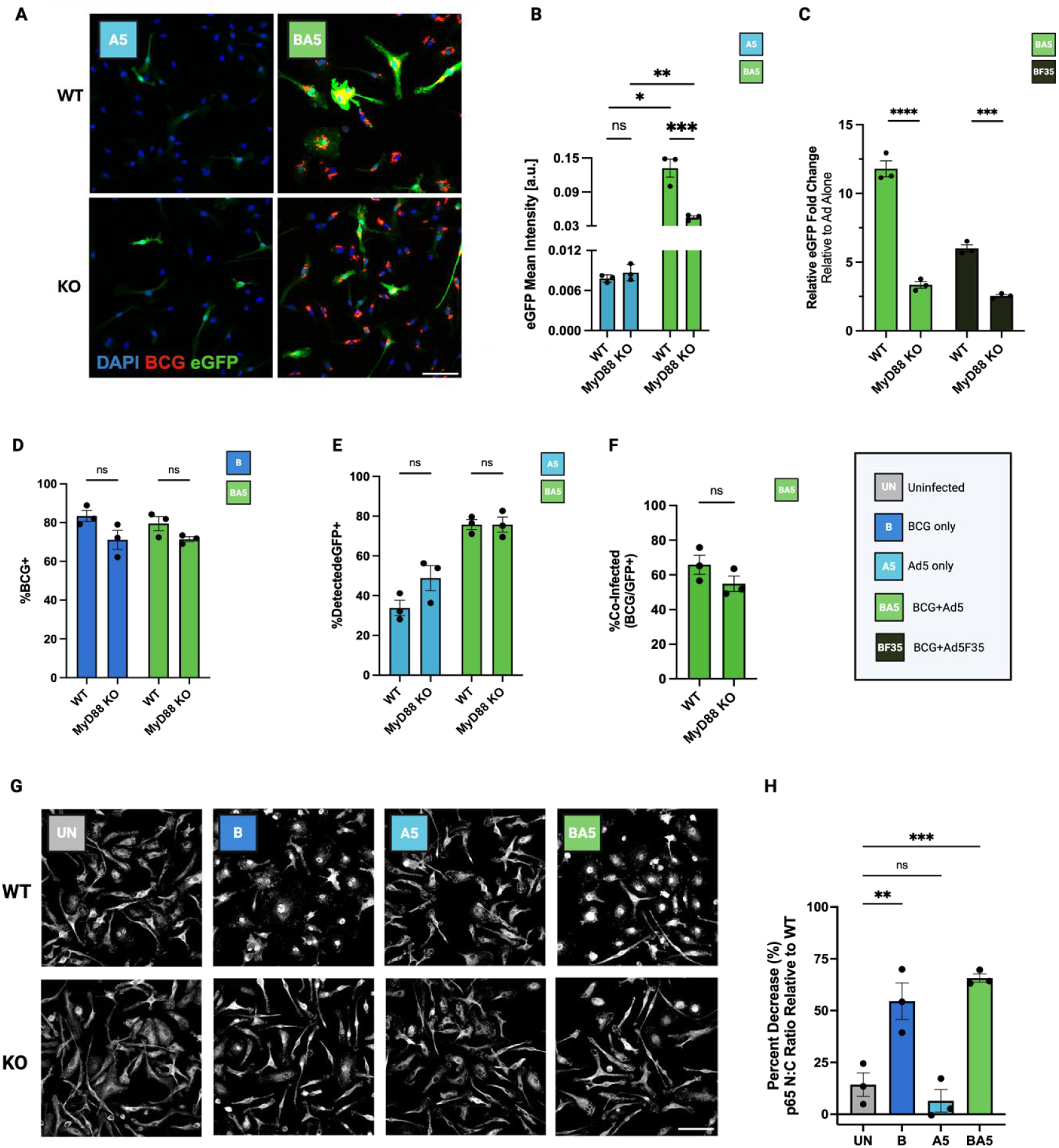
BCG drives higher adenovirus eGFP transgene expression in a MyD88-dependent manner. **(A)** WT and MyD88^-/-^ BMDMs were infected with BCG (MOI 10), and/or adenovirus (MOI 200). Representative images of eGFP expression in WT or MyD88^-/-^ BMDMs infected for 24 hours with Ad5, BCG or in combination. **(B)** eGFP Mean intensities among GFP-positive cells were quantified using a CellProfiler pipeline at 24 hpi. Two-way ANOVA with Sidak’s multiple comparisons test performed for significance. Unpaired t-test with Welch’s correction performed between A5 vs. BA5 for both WT and MyD88^-/-^ BMDMs. **(C)** RNA was isolated from wildtype and MyD88^-/-^ BMDM whole cell lysate at 4 hpi and analyzed for eGFP transcription. Fold changes in co-administered BMDMs were calculated using the 2^-ΔΔCT^ relative to their respective Ad5 or Ad5F35 alone groups. Two-way ANOVA with Sidak’s multiple comparisons test performed for significance. WT and MyD88^-/-^ BMDMs infected with **(D**) BCG or **(E)** Ad5 or **(F)** co-infected were quantified using a CellProfiler pipeline at 24 hpi. Two-way ANOVA with Sidak’s multiple comparisons test performed for significance for BCG, Ad5 infection rates. Unpaired t-test with Welch’s correction performed for co-infection rates. **(G)** Representative images of NF-κB/p65 staining in wildtype and MyD88^-/-^ BMDMs at 24 hours. Scale bars equal 50 µm. **(H)** Corresponding nuclear to cytoplasmic mean intensity ratios of NF-κB/p65 were calculated using a CellProfiler Pipeline. Outliers were identified and removed using the ROUT method with a maximum desired false discovery rate (Q) of 0.1%. Ratios of MyD88^-/-^ BMDMs reported as percentage decrease relative to WT across all cells. One-way ANOVA with Dunnett’s multiple comparisons test performed for significance. Data are presented as means ± SEM of three biological replicates (n=3). (ns p > 0.05, *p < 0.05, **p < 0.01, ***p < 0.001, ****p < 0.0001)

To confirm the effect of diminished eGFP expression in the MyD88^-/-^ BMDMs was not due to an impaired ability of the macrophages to internalize BCG or adenovirus, we quantified infection rates and observed no significant differences in uptake of either BCG or Ad5 between wildtype and MyD88^- /-^ BMDMs (**Figure 1D-F)**. Under these infection conditions, which were used in subsequent studies, we observed approximately 80% of the BMDMs infected with only BCG and 40% infected with Ad5 (**Figure 1D-E**). In BMDMs infected with both BCG and Ad5, the co-infection rate was approximately 60% (**Figure 1F**), indicating most of the BMDMs internalized both BCG and Ad5.

To determine if there was a link between elevated GFP expression and the transcription factor, NF-κB, we first performed p65 staining in wildtype and MyD88^-/-^ cells. As expected, in BMDMs lacking MyD88, there was a significantly reduced p65 nuclear to cytoplasmic ratio in BCG and co-infected cells (**Figure 1G-H)**. Staining for p65 in the BCG-infected and co-infected MyD88 knockout BMDMs resembled uninfected controls, with more diffuse, cytoplasmic staining when compared to the wildtype control (**Figure 1G**).

### Ad5, Ad5F35 infected macrophages induce STING-dependent type-I IFN production

To initially characterize the immune response to Ad5 and Ad5F35, we infected BMDMs with BCG and RdAds, either alone or in combination. Both RdAd- and BCG-infected BMDMs induced significantly higher *Ifnb1* transcription relative to uninfected at 4 hours post-infection (hpi), with adenoviruses inducing roughly 20-fold more *Ifnb1* transcription compared to BCG alone (**Figure 2A**). Co-infected macrophages induced comparable levels of *Ifnb1* transcription to RdAds alone, indicating that *Ifnb1* transcription is largely adenovirus driven at this timepoint (**Figure 2A**). BCG induced significantly higher *Tnf* transcription compared to uninfected BMDMs while administered RdAds alone induced minimal *Tnf* transcription. The combination of BCG and RdAds resulted in *Tnf* transcription similar to BCG alone indicating that the *Tnf* response is largely BCG-driven in co-infected macrophages (**Figure 2B**). To determine if the observed type-I IFN production was mediated through a MyD88-dependent pathway we infected MyD88-deficient BMDMs with RdAd, BCG or their combination and measured *Ifnb1* transcription. Results showed that RdAd-induced *Ifnb1* is independent of MyD88 whereas BCG-induced *Ifnb1* production was largely MyD88-dependent (**Figure 2C**). As expected, BMDMs lacking MyD88 also showed significantly reduced *Tnf* responses in both BCG- and MSM-infected cells (**Figure S1**). Previous studies have identified a role for cGAS-STING in driving type-I IFN production in adenovirus-infected immune and epithelial cells (38, 40). To evaluate the cGAS-STING pathway in BMDMs, we pretreated the cells independently with two STING inhibitors H-151 and SN-011. Both inhibitors led to a limited but significant lowering of *Ifnb1* transcription in *M. smegmatis* (MSM) infected BMDMs (**Figure 2D**). MSM is known to induce a type-I IFN response in macrophages through a STING-dependent pathway (14). In contrast, the addition of H-151 or SN-011 resulted in an almost complete loss of Ad5 and AdF35-induced *Ifnb1* transcription, indicating that STING is the major driver of type-I IFN production in these cells. This trend was qualitatively confirmed via immunocytochemistry with macrophages infected with adenovirus staining positive for perinuclear, Golgi-associated phospho-STING (**Figure 2E**).

**Figure 2.**
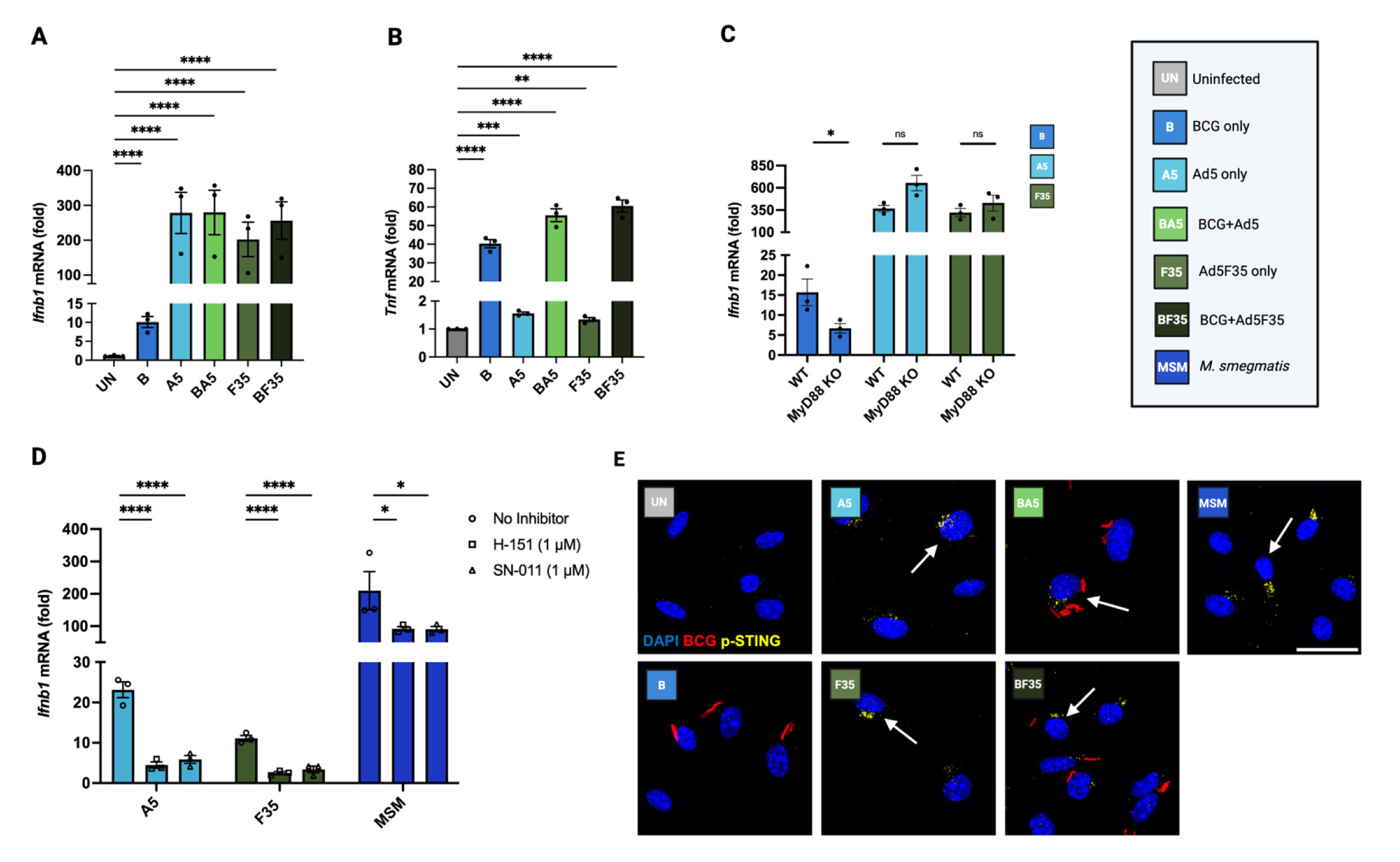
Ad5, Ad5F35 drive STING-dependent *Ifnb1* transcription in macrophages. BMDMs were infected with BCG (MOI 10), Ad5 or Ad5F35 (MOI 200), or in combination for 4 hours. RNA was isolated from whole cell lysate and analyzed by RT-qPCR for **(A)** *Ifnb1* and **(B)** *Tnf* transcript levels. One-way ANOVA with Dunnett’s multiple comparisons performed for significance. **(C)** *Ifnb1* transcription at 4 hpi in wildtype or MyD88^-/-^ BMDMs infected with BCG (MOI 10) or adenovirus (MOI 200) and analyzed by RT-qPCR. Two-way ANOVA with Sidak’s multiple comparisons test performed for significance. **(D)** *Ifnb1* transcription in BMDMs pretreated with STING inhibitors H-151 and SN-011 at 1 µM for 3 hours prior to a 4 hour infection with adenovirus (MOI 50) and 8 hour infection for MSM (MOI 10) analyzed by RT-qPCR. Two-way ANOVA with Dunnett’s multiple comparisons test performed for significance. Data are presented as means ± standard error of mean (± SEM) of three biological replicates (n=3). (ns p > 0.05, *p < 0.05, **p < 0.01, ***p < 0.001, ****p < 0.0001) **(E)** Representative images of phospho-STING positive BMDMs from five independent fields indicated by white arrows. Scale bar equals 25 µm.

### BCG-mediated suppression of the type-I IFN response in BCG+Ad5 co-infected macrophages

To understand the dynamics of *Ifnb1* transcription over time, we isolated RNA at various times post-infection ranging from 1.5 to 12 hpi. The time course studies revealed that Ad5 induced significantly higher *Ifnb1* transcription compared to BCG-infected BMDMs at multiple time points (**Figure 3A).** Ad5-induced *Ifnb1* transcription reached optimal transcription at around 5 hpi with BMDMs infected with adenovirus alone exhibiting significantly higher *Ifnb1* transcription compared to co-infected BMDMs. This observed suppression by BCG was most evident at 5 hpi but was observed at multiple time points (**Figure 3A**). To determine if this effect was also observed at the protein level, we measured IFN-β in cell culture supernatants at 8 hpi. There was significantly higher IFN-β in BMDMs infected with Ad5 alone compared to co-infected cells (**Figure 3B**). At this time point there was no significant difference in IFN-β levels between BCG-infected and uninfected cells (**Figure 3B**). These data suggest limited autocrine or paracrine signaling in co-infected cells compared to Ad5-infection alone. To understand the autocrine and paracrine signaling dynamics, we measured the phosphorylation state of STAT1 and expression of IRF7. Phospho-STAT1 levels were elevated by 5 hpi and remained elevated in Ad5-infected macrophages while BCG-induced STAT1 activation began later between 6.5 and 8 hpi (**Figure 3C**). When co-infected with BCG and Ad5, BMDMs exhibited delayed and suppressed STAT1 activation compared to those treated with Ad5 alone, with densitometry revealing significantly lower levels of phospho-STAT1 from 5 through 10 hpi compared to Ad5 only infected group (**Figure 3D**). When compared to the BCG-alone, STAT1 activation in the co-infected BMMs was marginally higher (**Figure 3D**). Interestingly, phospho-STAT1 activated in response to BCG failed to stimulate the increased expression of the type-I IFN transcription factor IRF7 (**Figure 3C**). IRF7 expression was observed in the co-infected BMMs but the production was less than in Ad5-infected BMMs (**Figure 3E**). These results indicate a temporal dynamics to adenovirus-induced STAT1 phosphorylation and IRF7 expression, but the magnitude of this response is suppressed by BCG when added in combination. To determine if BCG co-infection with Ad5 limits the normally viral-induced expression of ISGs we measured the transcription of two canonical ISGs: *Cxcl10* (**Figure 3F**) and *Ifit2* (**Figure 3G**). Upon Ad5 infection, transcription for both ISGs began at 5 hpi, peaked at 8 hpi, and began returning to baseline by 12 hpi. The delayed induction corresponds to the time required for IFN-β production and subsequent binding to the IFNAR and activation of the JAK/STAT pathway. The adenovirus infection alone induced roughly 10-fold higher ISG transcription than the co-infected group indicating that BCG infection suppressed ISG expression likely due, in part, to decreased IFN-β produced in co-infected BMMs compared to Ad5 infected cells **(Figure 3F-G)**. Other mechanisms that could account for the approximately 90% decrease in *Cxcl10* and *lfit2* expression in co-infected BMMs could stem from factors that are known to block signaling initiated from IFNAR. *Socs1* is a ISG with a well-established role in inhibiting type-I IFN-induced STAT1 activation (8). At 4 hpi, *Socs1* transcript levels were significantly increased in co-infected macrophages compared to each component alone (**Figure 3H**). This additive effect on *Socs1* transcription was predominantly mediated by BCG-induced *Socs1,* with BCG infection leading to significantly higher *Socs1* transcription compared to Ad5 alone (**Figure 3H**). We also analyzed transcript levels of *Usp18*, an ISG that directly inhibits paracrine JAK-STAT signaling, but observed no significant differences among groups compared to uninfected at 4 hpi, indicating this effector is likely not a major driver for the observed suppression (**Figure S2**).

**Figure 3.**
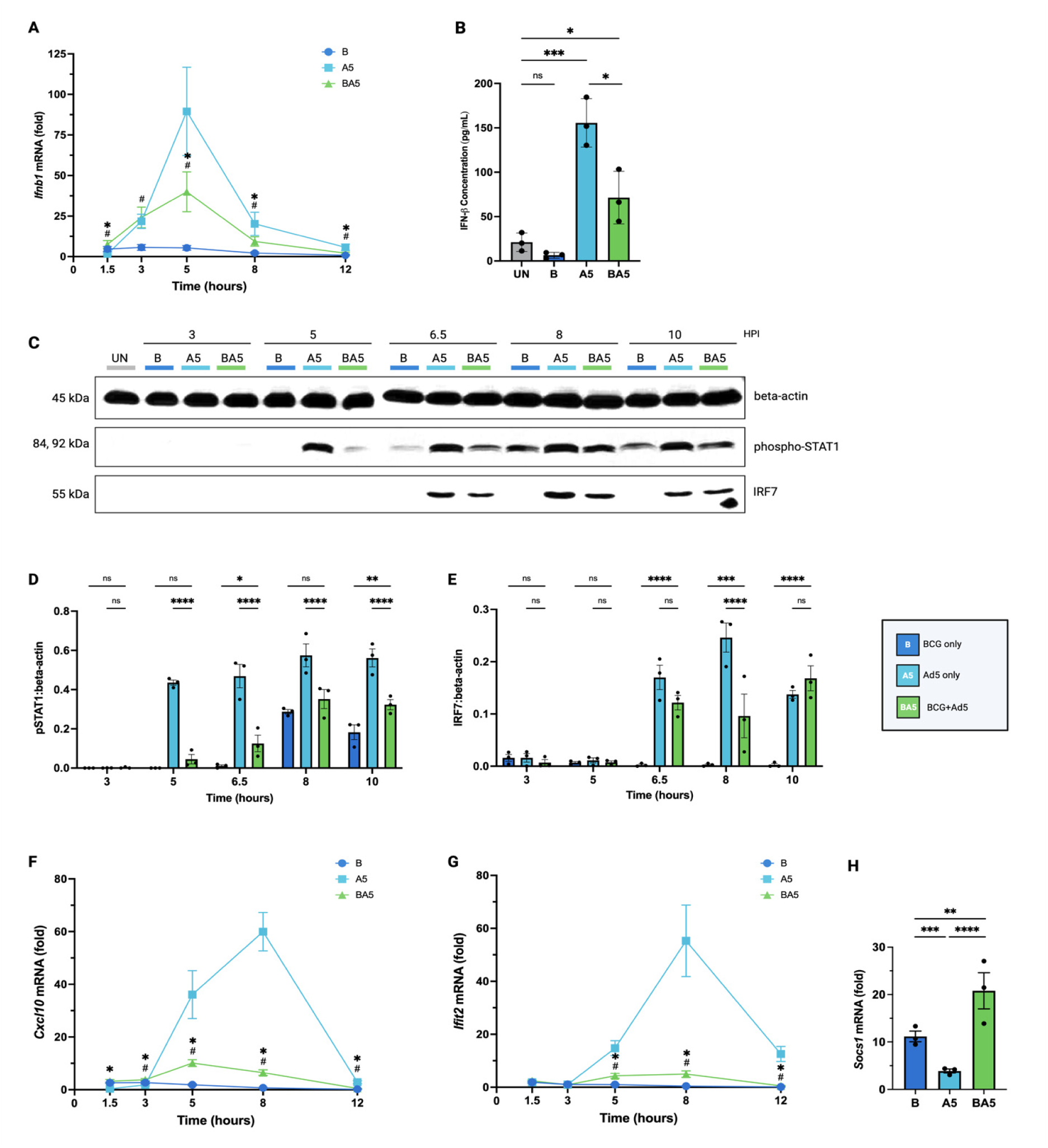
BCG suppresses adenovirus-driven type-I IFN responses in co-infected macrophages. BMDMs were infected with BCG (MOI 10), Ad5 (MOI 200), or in combination. **(A)** RNA was isolated from whole cell lysate at various hours post-infection and analyzed for *Ifnb1* transcription by RT-qPCR. **(B)** Secreted IFN-β levels in BMDM cell culture supernatant measured at 8 hpi via ProQuantum Immunoassay. One-way ANOVA with Dunnett’s multiple comparisons performed for significance. Unpaired t-test with Welch’s correction performed between A5 vs. BA5. Values below the limit of detection were assigned LOD/2. **(C)** Western blot showing phospho-STAT and IRF7 levels at various times post infection in whole cell lysate. Beta-actin served as a loading control. Densitometry was calculated as ratios for **(D)** phospho-STAT1 and **(E)** IRF7 relative to beta-actin using ImageJ. Two-way ANOVA with Dunnett’s multiple comparisons test was performed for significance. RNA was isolated from whole cell lysate at various hours post-infection and analyzed for **(F)** *Cxcl10* and **(G)** *Ifit2* transcription by RT-qPCR. **(H)** RNA was isolated from whole cell lysate at 4 hpi and analyzed for *Socs1* transcription by RT-qPCR. One-way ANOVA performed for significance with Sidak’s multiple comparisons. Fold changes were calculated using the 2^-ΔΔCT^ relative to uninfected BMDMs. (ns p > 0.05, *p < 0.05, **p < 0.01, ***p < 0.001, ****p < 0.0001) For time course RT-qPCR, two-way ANOVA with Dunnett’s multiple comparisons test was performed for significance. (#p < 0.05 for B vs. BA5, *p < 0.05 for A5 vs. BA5) Data are presented as means ± standard error of mean (± SEM) of three biological replicates (n=3).

### BCG infection drives pro-inflammatory responses in co-infected macrophages

To investigate whether adenovirus co-infection influences the pro-inflammatory response induced by BCG we initially focused on TNF, a well-established cytokine associated with macrophage activation. To understand the dynamics, we analyzed *Tnf* transcription with RNA isolated at different times post-infection with BCG, Ad5, or in combination. Results indicated *Tnf* responses were largely BCG-driven with no significant differences between co-infected BMDMs (**Figure 4A**). *Tnf* transcription in BCG- and co- administered macrophages peaked early after infection and began to decrease by 8 hpi (**Figure 4A**). Adenovirus infection alone induced minimal *Tnf* transcription. To look at signaling pathways known to be associated with pro-inflammatory responses including production of TNF, we performed immunoblotting of phosphorylated-p38 MAPK. Similarly, we observed higher levels of p38 MAPK activation in BCG- and co-infected BMDMs and minimal induction by Ad5 alone (**Figure 4B**). Co-infected BMDMs induced similar levels of p38 activation compared to macrophages infected with BCG alone (**Figure 4C**). NF-kB is a transcription factor associated with production of many pro-inflammatory mediators, including TNF. To look at the activation of NF-kB following the different infections, we performed immunofluorescent staining via confocal microscopy to measure NF-kB p65 translocation to the nucleus. Quantification of images using a Cell Profiler pipeline revealed significantly higher nuclear to cytoplasmic ratios of p65 mean intensities for BCG and co-infected BMDMs compared to uninfected cells (**Figure 4D-E**). No difference in p65 activation was observed between Ad5-infected and uninfected BMMs (**Figure 4E**), further indicating the adenovirus is not a major driver of pro-inflammatory pathways in macrophages under these experimental conditions.

**Figure 4.**
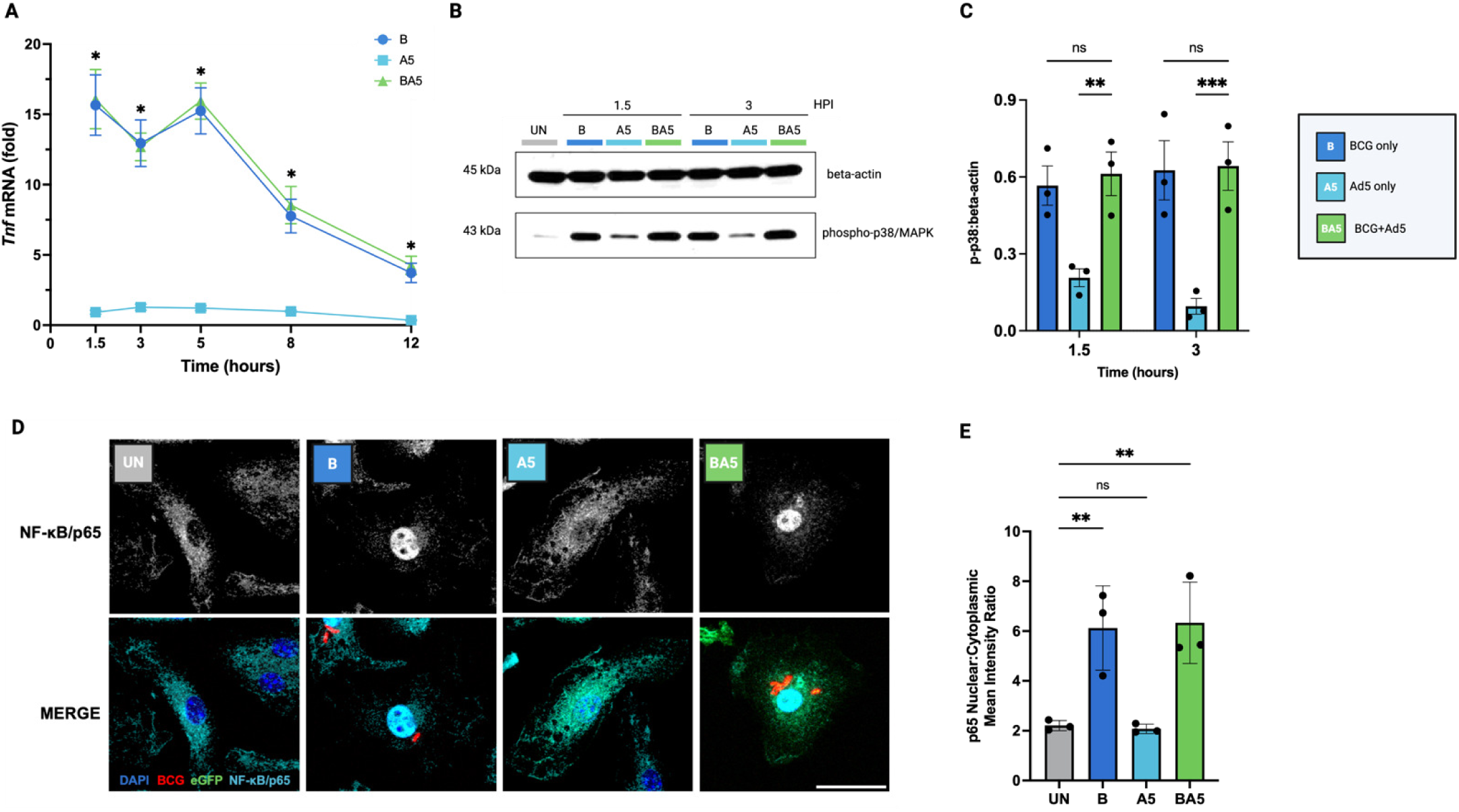
BCG drives pro-inflammatory TNF-α signaling pathways in co-infected macrophages. **(A)** BMDMs were infected with BCG (MOI 10), Ad5 (MOI 200), or in combination. RNA was isolated from whole cell lysate at various hours post-infection and analyzed for *Tnf* transcription by RT-qPCR. Two-way ANOVA with Dunnett’s multiple comparisons test was performed for significance. (# p < 0.05 for B vs. BA5, * p < 0.05 for A5 vs. BA5) **(B)** Western blot showing phospho-p38/MAPK levels at 1.5 and 3 hpi. Beta-actin served as a loading control. **(C)** Densitometry was calculated as ratios for phospho-p38/MAPK relative to beta-actin using Image J. Two-way ANOVA with Dunnett’s multiple comparisons test was performed for significance. **(D)** Representative images of NF-κB/p65 staining in BMDMs. Scale bar equals 25 µm. **(E)** Nuclear to cytoplasmic mean intensity ratios of NF-κB/p65 were calculated using a CellProfiler pipeline across all cells. Outliers were identified and removed using the ROUT method with a maximum desired false discovery rate (Q) of 0.1%. One-way ANOVA with Dunnett’s multiple comparisons test performed for significance. Data are presented as means ± standard error of mean (± SEM) of three biological replicates (n=3). (ns p > 0.05, **p < 0.01, ***p < 0.001).

### Macrophages co-infected with BCG and Ad5 lead to additive expression of host protective genes

In addition to TNF, we also looked at *Ccl5, Acod1,* and *Nos2* expression due to their established host-protective roles during an MTB infection. In contrast to ISGs, time course experiments revealed late, additive trends in co-infected BMDMs for these three genes (**Figures 5A-C**). Co-infected BMDMs induced significantly higher *Ccl5* transcription when compared to each component alone at both 8 and 12 hpi (**Figure 5A**). *Acod1* transcription followed a similar trend with a modest additive effect by 8 and 12 hpi in co-infected BMDMs (**Figure 5B**). *Nos2* transcription was strongly induced in BCG-infected BMDMs, over 300 fold by 3 hpi compared to uninfected cells. Co-infected macrophages showed a similar trend (**Figure 5C**). Ad5-induced *Nos2* transcription began at 5 hpi and peaked around 8 hpi, which at this timepoint showed similar expression to BCG infected BMDMs. At the 8 and 12 hour time point there appeared to be an additive effect in co-infected BMDMs as the *Nos2* levels were more than BCG and Ad5 alone, although the differences did not reach statistical significance (**Figure 5C**). We performed western blotting to confirm this additive trend by probing for iNOS expression at 5, 8, and 12 hpi (**Figure 5D**). Immunoblotting and densitometry revealed significantly higher iNOS protein expression at 8 hpi in co-infected macrophages compared to BMMs infected with BCG or adenovirus alone (**Figure 5D-E**). Although higher expression in co-infected BMMs was also observed at 12 hpi, this effect did not reach statistical significance compared to BCG-induced iNOS expression (**Figure 5E**).

**Figure 5.**
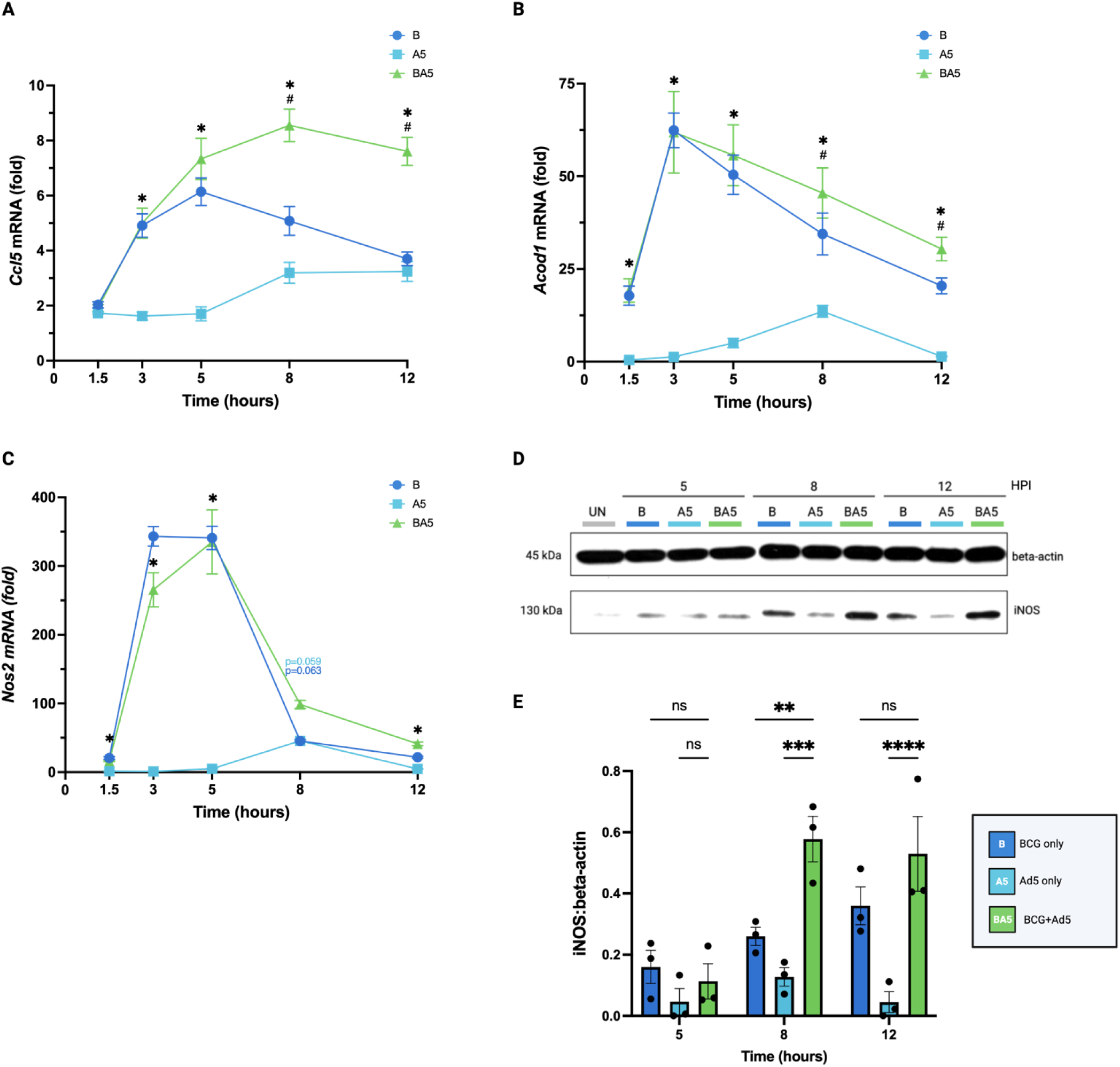
Co-infection with BCG and Ad5 elicits late, additive expression of Ccl5, Acod1 and iNOS in macrophages. BMDMs were infected with BCG (MOI 10), Ad5 (MOI 200), or in combination. RNA was isolated from whole cell lysate at various hours post-infection and analyzed for **(A)** *Ccl5* **(B)** *Acod1, and* **(C)** *Nos2* transcription by RT-qPCR. Two-way ANOVA with Dunnett’s multiple comparisons test was performed for significance. (# p < 0.05 for B vs. BA5, * p < 0.05 for A vs. BA5) **(D)** Western blot of iNOS expression levels at 5, 8, and 12 hpi. Beta-actin served as a loading control. **(E)** Densitometry was calculated as ratios for iNOS relative to beta-actin using ImageJ. Two-way ANOVA with Dunnett’s multiple comparisons test was performed for significance. Data are presented as means ± standard error of mean (± SEM) of three biological replicates (n=3). (ns p > 0.05, *p < 0.05, **p < 0.01, ***p < 0.001, ****p < 0.0001

## DISCUSSION

Recombinant adenoviruses have been used previously as a mechanism to boost BCG-mediated vaccine efficacy. The general protocol entails BCG vaccination followed months to years later by the administration of RdAds carrying DNA encoding one or more mycobacterial antigens (41–46). Unlike these conventional BCG prime and boost regimens, we aimed to modulate innate immunity through their simultaneous administration. However, since both adenovirus and BCG can infect macrophages, it is important to define how the co-infection would affect the macrophage response, especially in the context of Type-I IFN and pro-inflammatory responses. Here, we demonstrate that RdAds can serve as potent immunomodulators in primary macrophages when combined with *M. bovis* BCG. Notably, this co-administration reveals a complex regulatory interplay wherein BCG works to dampen adenovirus-triggered type-I IFN signaling pathways yet drives additive expression in genes with established protective roles in MTB infection. Our results also demonstrate how BCG can drive higher adenovirus transgene expression during co-administration through a mechanism dependent on MyD88.

Ad5 and Ad5F35 have been shown to utilize receptors coxsackie and adenovirus receptor (CAR) and CD46, a human complement regulatory receptor, respectively (47, 48). In cells expressing low or absent levels of their receptors, such as murine macrophages, these adenoviruses can be internalized via interactions with surface integrins, Fc, complement, and scavenger receptors (49, 50). In this study, we demonstrate both adenovirus serotypes were able to infect BMDMs alone and when co-administered with BCG. Although showing delayed kinetics of transduction in BMDMs (49), the RdAds were able to induce robust type-I IFN responses, as reported previously (38, 40, 51), in a cGAS-STING-dependent and MyD88-independent manner (51, 52). Contrary to type-I IFN responses, Ad5 and Ad5F35 were not major drivers of a TNF-α response and induced minimal to modest activation of various signaling pathways that lead to TNF-α, such as p38 MAPK (52). Interestingly, we noted enhanced transgene transcription in BCG and adenovirus co-infected macrophages. This enhancement of eGFP was attenuated in MyD88 knockout BMDMs, a trend that correlated with decreased nuclear NF-κB subunit p65 in BCG-infected and co-infected BMDMs. We hypothesize this effect is driven by the binding of BCG-induced transcription factors on the CMV promoter. Binding sites on the CMV promoter for certain transcription factors have been described previously (53, 54).

Since knocking out MyD88 abolishes nearly all p65 nuclear translocation but fails to return eGFP levels to baseline, it suggests that p65 is not the sole driving factor for this observed effect. Other transcription factors induced by BCG infection, such as AP-1 and ATF/CREB, could also be involved as they have been shown to influence CMV promoter activity (54, 55). This observation could have important implications for heterologous prime-boost strategies since macrophages infected with BCG will also promote higher expression of adenovirus-delivered transgene, which could influence the elicited immune response.

Given the importance of bolstering type-I IFN in the context of BCG vaccination, we worked to further characterize the type-I IFN signaling pathways in macrophages upon BCG and Ad5 co-infection. While early *Ifnb1* transcription levels mirrored those observed in adenovirus only infected BMDMs, we observed a suppression of transcription and secretion of IFN-β in co-infected macrophages. This coincided with the start of adenovirus-induced autocrine or paracrine Type-I IFN signaling as measured by phosphorylation of STAT1 and correlated with a BCG-mediated suppression of STAT1 activation in co-infected BMDMs. Both BCG- and MTB-infected BMDMs were previously shown to suppress STAT1 phosphorylation (56). In MTB-infected cells, the suppression was demonstrated to be specific to type-I IFNs and due to reduced phosphorylation of INFAR-associated JAK1 and TYK2, which resulted in decreased phosphorylation levels of STAT1 and STAT2 (56). These findings align with our observations, providing a possible mechanistic explanation for the BCG-mediated suppression of STAT1 activation seen in our model; however, further studies are needed to test this hypothesis. We also demonstrated that co-infected macrophages can induce higher, earlier expression of SOCS1, which is an established inhibitor of JAK-mediated STAT1 phosphorylation (34, 57, 58).

Both BCG and Ad5 infection lead to STAT1 activation. The mechanism by which BCG induces STAT1 phosphorylation is unclear since it induces only a weak Type-I IFN response. It is known that BCG induces IL-6 and IL-10 expression (60, 61), which can lead to STAT3 activation and, to a lesser extent, STAT1 phosphorylation (62, 63). Hence, the late STAT1 phosphorylation observed in BCG-infected BMDMs could be a result from a pro-inflammatory paracrine cytokine signaling independent of IFNAR. Interestingly, despite BCG-induced STAT1 phosphorylation, only BMDMs infected with Ad5, either alone or in combination with BCG, led to increased expression of ISGs. In addition to the type-I IFN transcription factor IRF7, we analyzed CXCL10 and IFIT2, two canonical ISGs with protective roles in the context of MTB infection. CXCL10, C-X-C motif chemokine ligand 10, is a crucial inflammatory chemokine that attracts immune cells to the site of TB infection and facilitates granuloma formation (64) whereas IFIT2, Interferon-Induced Protein with Tetratricopeptide Repeats 2, is an innate immune effector that has been shown to be involved in the macrophage response to MTB infection and restrict mycobacterial growth (65, 66). Similar to IFN-β expression and STAT1 activation, co-infected BMDMs exhibited significantly blunted ISG responses compared to Ad5 alone. A similar blunting effect on ISGs has been observed in the context of MTB infection in BMDMs (66). In addition to inhibiting pathways that lead to the activation of STAT1, BCG could inhibit the formation of ISGF3 downstream of STAT1. Proteins like PIAS1 (protein inhibitor of activated STAT1) that block STAT1 DNA-binding activity and abrogate the ISG response could be induced by BCG infection, especially since PIAS can be activated by pro-inflammatory stimuli like TNF-α and lipopolysaccharide (LPS) (8, 67). This is also a potential mechanism to explain why, despite STAT1 phosphorylation in BCG-infected macrophages, no ISG expression was observed. However, this hypothesis and whether it applies to MTB, requires further investigation.

There is growing evidence that an MTB infection followed by a subsequent viral challenge leads to accelerated bacterial progression and compromised host survival, driven largely by detrimental type-I IFN signaling (68, 69). Our data with BCG and published data with MTB (66) which shows that mycobacteria can suppress type-I IFN signaling in macrophages, suggest a potential impaired ability to respond to viral pathogens in TB patients (70). A failure of macrophages infected with mycobacteria to properly mount a type-I IFN response during the acute phase of viral infection, could lead to an environment more permissive to viral replication.

Unlike the antagonism observed in ISG expression, we highlighted three genes in which BCG and Ad5 co-infection led to additive effects on expression: CCL5, ACOD1, and iNOS. CCL5, also known as RANTES, is a well-established chemokine that plays protective roles in granuloma formation in MTB infection (71) whereas ACOD1, also known as IRG1, and iNOS are metabolic enzymes that are involved in restricting intracellular growth of MTB (72–76). The additive effects of co-administering BCG with Ad5 were observed following the onset of autocrine and paracrine signaling. The differences between the suppressed ISG expression and enhanced, additive effects seen with these genes is likely due to their differences in the transcription factors involved. Unlike ISGs which primarily depend on ISGF3 to promote expression, the genes in which additive effects were observed can be activated by various transcription factors, including STAT1-homodimers binding GAS elements, Interferon Regulatory Factors (IRFs), AP-1, and/or or NF-κB (77–80). We hypothesize that this additive trend in co-administered macrophages results from enhanceosome formation, with BCG and Ad5 providing complementary transcription factors that cooperatively drive transcription to induce higher levels of expression achieved by either component individually.

Together, our results show that BCG and Ad5 co-infection result in a unique gene expression profile when compared to infections with BCG or adenovirus alone. This varied response by the BMDMs to co-infection likely stems from its differential effect on the activation of STAT1, NF-κB and likely other transcription factors. The administration of RdAds simultaneously with BCG presents a previously unexplored way to modulate macrophage signaling and may offer alternative methods to bolster BCG-conferred protection.

## METHODS

### Mycobacterial Cultures and Adenoviral Stocks

*M. bovis* BCG:pMV261-DsRed was generated as described previously (81). Mycobacteria were grown in Middlebrook 7H9 broth medium (Difco, Becton-Dickinson) containing 10% OADC [oleic acid (Thermo), dextrose (Sigma), albumin (Sigma Aldrich), and catalase(Thermo)], 0.05% Tween 80 (Sigma), and 50 µg/mL hygromycin B (Sigma) for DsRed BCG until exponential phase, aliquoted, and stored in cryovials (Corning) at −80 °C until use. Prior to use, the DsRed BCG was de-clumped by syringe-passage with a 27-gauge blunt-end needle (Thermo, BD) at least 5 times. Mycobacterial culture aliquots were thawed and plated on Middlebrook’s 7H10 (Sigma) agar plates supplemented with 10% OADC and 50 µg/mL hygromycin B for DsRed BCG. Colonies stored at 37°C and enumerated 2 to 4 days later for *M. smegmatis* and 2 to 3 weeks later for DsRed BCG. Adenovirus stocks (Ad5-CMV-eGFP, Ad5F35-CMV-eGFP) were supplied by the Gene Vector Core at Baylor College of Medicine, and internally confirmed to be replication deficient as well as titered via plaque-forming assay.

### Mice

Wild type and MyD88^-/-^ C57BL/6 mice (Jackson Laboratory) were housed in the Freimann Life Science Center at the University of Notre Dame. All animals were cared for in accordance with the Association for Assessment and Accreditation of Laboratory Animal Care International under pathogen-free conditions, and all animal experiments were approved by the University of Notre Dame’s Institutional Animal Care and Use Committee.

### Mammalian cell culture

Bone marrow derived macrophages (BMDMs) were isolated from the femurs of 6- to 8-week-old C57BL/6 mice (Jackson Laboratory), maintained in BMDM media (10% HI-FBS, 20% L929, 1% penicillin-streptomycin), and processed as previously described (82).

### Macrophage infections

Bone marrow derived macrophages (BMDMs) were cultured in BMDM media for seven days on non-tissue culture plates, with media changed at 24 and 96 hours. On day seven, macrophage monolayers were washed with sterile PBS (Cytiva) and dissociated with 0.05% Trypsin EDTA in PBS. Trypsin was inactivated with the addition of BMDM media and the cells were centrifuged at 1000 rpm for 5 minutes. Cells were then resuspended in fresh BMDM media, enumerated using a hemocytometer, and seeded into wells of tissue-culture treated plates (Greiner Bio-One). Cells were seeded at a density of 1.5 x10^5^ to 2.5 x10^5^ in 12 well plates and 4 x10^5^ to 5 x10^5^ in 6 well-plates. Cells were incubated overnight at 37°C with 5% CO2. The following day, macrophages were washed and replaced with antibiotic-free media. Cells were then infected at an MOI of 10 for bacteria and 200 for adenovirus unless indicated otherwise. After 4 hours, media was removed and cells washed with BMDM media to remove extracellular bacteria and virus. BMDM media was added back to the cells and samples were collected at various times post infection. For STING inhibitor experiments, BMDMs were treated with 1 µM of H-151 or SN-011 (MedChem Express) for three hours prior to infection, and RNA was collected at 4 hours post infection.

### Western blotting

Whole cell lysates were generated using RIPA buffer (Sigma) supplemented Halt protease and phosphatase inhibitor (Thermo). The resulting whole cell lysates were centrifuged at 12,000 rpm for 2 minutes before quantifying protein concentration using a Pierce BCA Protein Assay Kit (Thermo). Equal amounts of protein were loaded into lanes of 10-12% SDS-PAGE gels (Bio-Rad). Following electrophoresis, the separated proteins were transferred onto a methanol activated 0.45 μM PVDF membrane (Sigma) at 20V overnight at 4°C in Transfer Buffer (192 mM Glycine, 25 mM Tris Base, and 10% Methanol). Membranes were blocked in 5% bovine serum albumin fraction V (BSA; Thermo) diluted in TBS-T (20 mM Tris Base, 150 mM NaCl, and 0.5% tween 20; pH = 7.6) for one hour at room temperature with gentle agitation. Membranes were then incubated with primary antibody diluted in 5% BSA in TBS-T overnight rocking at 4°C. The following day, membranes were washed 5 times for 5 minutes each in TBS-T before incubation with an HRP-conjugated secondary Goat anti-Rabbit IgG antibody (Thermo) diluted in 5% BSA in TBS-T at room temperature for 45 minutes. Membranes were washed 5 times in TBS-T for 5 minutes. Bound antibody were detected using SuperSignal West PicoPlus Chemiluminescent or Femto Maximum Sensitivity Substrate (Thermo), exposed to BioBlue-MR Autoradiography film (Alkali Scientific), and imaged on a Konica SRX-101A developer. See **Table S1** for a list of antibodies utilized in this study. Densitometric analysis of western blots was performed using ImageJ. Target bands were normalized to the signal of the corresponding loading control (β-actin).

### ProQuantum Immunoassay

Cell culture supernatants were collected, spin-clarified at 12,000 rpm for 90 seconds, aliquoted, and frozen at −80°C until use. Samples were analyzed using the Mouse IFN beta ProQuantum Immunoassay Kit (Thermo) according to the manufacturer’s recommendations.

### Reverse transcription quantitative PCR

RNA from BMDM monolayers were isolated using Quick-RNA Miniprep Kit (Zymo). Cells were directly lysed in wells of a 12-well plate in RNA lysis buffer buffer and DNase treated and processed as per the manufacturer’s recommendation. The isolated RNA was assessed for quality and concentration by absorbance at 260nm using a Nanodrop 2000 Spectrophotometer (Thermo). For quantitative PCR analysis, 20-50 ng of isolated RNA was utilized as input on a QuantStudio 5 Real-Time PCR System (Thermo) using the the Luna^®^ Universal One-Step RT-qPCR Kit (New England Biolabs). Raw Ct values were quantified using the ΔΔCt method. The primer pairs used in this study are listed in **Table S2**.

### Immunocytochemistry

Cells were fixed at various time points post infection with ice-cold PFA (Sigma) for 15 minutes, before three subsequent washes in PBS. Cells were then permeabilized with PBS containing 0.1% Triton X-100 (Sigma) for 5 minutes before washing in PBS and blocking with 3% BSA in PBS-Triton for 30 minutes at room temperature. Coverslips (Thermo) were transferred onto 125 µl droplets of primary antibody and incubated overnight at 4°C (phospho-STING) or 2 hours at room temperature (NF-kB/p65). Cells were washed once with PBS, followed by PBS-TritonX, and another PBS wash. Cells were incubated with Alexa Fluor 647 secondary antibody (Thermo) for 1 hour at room temperature. Coverslips were washed three times as described previously and mounted with one drop of Fluoromount-G Mounting Medium with DAPI (Thermo). Coverslips were imaged using Leica Stellaris 8 DIVE located at the Notre Dame Integrated Imaging Facility. Mean intensities were calculated using a CellProfiler pipeline. **Table S1** lists the antibodies used in this study.

## Acknowledgements

We are grateful for the technical expertise and guidance of members of the University of Notre Dame Integrated Imaging Facility, especially Dr. Sara Cole. We also thank the Gene Vector Core at the Baylor College of Medicine for supplying the adenovirus stocks, specifically Kazuhiro Oka, Ph.D. (Director), Corinne Sonnet, Ph.D. (Co-Director) as well as Sean Cuellar (Staff).

## Author Contributions

J.V. and J.S.S. contributed to conceptualization and methodology. J.V. carried out the experiments, analyses, and data visualization. J.V. prepared the original draft of the manuscript and carried out revisions under the critical guidance of J.S.S. J.S.S. was responsible for supervision, project administration, and funding acquisition. All authors have read and agreed to the published version of the manuscript.

